# Hippo-*vgll3* signaling may contribute to sex differences in Atlantic salmon maturation age via contrasting adipose dynamics

**DOI:** 10.1101/2024.10.21.619349

**Authors:** Ehsan Pashay Ahi, Jukka-Pekka Verta, Johanna Kurko, Annukka Ruokolainen, Paul Vincent Debes, Craig R. Primmer

## Abstract

Sexual maturation in Atlantic salmon entails a transition in energy utilization, governed in part by genes and environmental stimuli in sex-specific manner. Salmon males require less energy, in the form of adiposity, to mature compared to females and typically mature younger. Maturation age is also influenced in a sex-dependent fashion by the *vgll3* genotype (*vestigial-like 3*), a co-factor in the Hippo pathway. The underlying molecular processes of sex-dependent maturation age, and how they interplay with adiposity and *vgll3* genotypes, have remained unknown. To elucidate the molecular mechanisms underlying sex- and genotype-specific maturation differences, we investigated the association of *early* (E) and *late* (L) maturation *vgll3* alleles and the transcriptional expression of > 330 genes involved in the regulation of the Hippo pathway and sexual maturation, and related to molecular signals in brain, adipose tissue, and gonads. The strongest effect of *vgll3* genotype was observed in adipose tissue for females and in brain for males, highlighting a sex-specific expression difference in the main association of *vgll3* genotype. Genes related to ovarian development showed increased expression in *vgll3*EE* compared to *vgll3*LL* females. Additionally, *vgll3*EE* females compared to *vgll3*EE* males exhibited reduced markers of pre-adipocyte differentiation and lipolysis yet enhanced expression of genes related to adipocyte maturation and lipid storage. Brain gene expression further showed sex-specific expression signals for genes related to hormones and lipids, as well as tight junction assembly. Overall, these sex-specific patterns point towards a greater lipid storage and slower energy utilization in females compared to males. These results suggest that Hippo-dependent mechanisms may be important mediators of sex differences in maturation age in salmon.

## Introduction

The timing of sexual maturation can have dramatic effects on survival and reproductive success (Mobley et al. 2021) and is influenced by environmental cues and genetic mechanisms (Taranger et al. 2010). Maturation age often exhibits sex-specific differences within species (Worthman & Trang 2018; Tarka et al. 2018). In Atlantic salmon, sexual maturation is linked with seasonal environmental changes with differences between sexes (Mobley et al. 2021). Lipid allocation variation also plays a role in determining maturation age, as individuals must accumulate sufficient energy reserves to initiate maturation (Rowe et al. 2011; House, Debes, Kurko, et al. 2023; Mogensen & Post 2012). Genetic factors, notably the vestigial-like family member 3 (*vgll3*) gene, serve as major determinants of maturation age with sex-specific effects in this species (Barson et al. 2015; Czorlich et al. 2018; Miettinen et al. 2023). *Vgll3* influences maturation timing in males as early as < 1 year of age in controlled conditions (Sinclair-Waters et al. 2021; Debes et al. 2021; Verta et al. 2020). Interestingly, associations with similar traits (pubertal timing, pubertal growth spurt) have been detected with the human ortholog (*VGLL3*) (Cousminer et al. 2016; Perry et al. 2014; Tu et al. 2015) and *VGLL3* has also been identified as a promoter of sex-biased autoimmune diseases (Liang et al. 2016).

Previous molecular studies of *vgll3* in Atlantic salmon have primarily focused on males for practical reasons such as the convenience of some males maturing as early as < 1 year of age, (Verta et al. 2024; Kurko et al. 2020; Verta et al. 2020; Ahi et al. 2022; Ahi, Frapin, et al. 2024; Ahi et al. 2023; Ahi, Verta, et al. 2024b), whereas females usually take three or more years to mature (Åsheim et al. 2023). Therefore, the underlying molecular processes of sex-specific maturation patterns have remained largely unexplored. Recent gene co-expression network analyses have indicated that the effects of *vgll3* genotypes in males on genes playing a role in the reproductive axis, adipogenesis and neurogenesis are exerted through the modulation of various components of the Hippo signaling pathway (Ahi et al. 2023; Ahi, Verta, et al. 2024b, 2024a). The Hippo pathway is known for its role in biological functions such as controlling organ size in vertebrates (Kjærner-Semb et al. 2018; Kurko et al. 2020; Sen Sharma et al. 2019), regulating adipocyte proliferation and differentiation (Ardestani et al. 2018; Halperin et al. 2013) and high fat diet-induced neural differentiation (Poon et al. 2013). In mammals, for instance, YAP1, a major transcription co-factor of the Hippo pathway, is required for adipogenesis (Ardestani et al. 2018), whereas VGLL3 functions as an inhibitor of adipogenesis (Halperin et al. 2013), indicating the significance of this pathway in energy acquisition processes. VGLL3 is thought to compete with YAP1, which acts as an inhibitor of the Hippo pathway (Hori et al. 2020; Kurko et al. 2020). Moreover, the Hippo pathway has been implicated in responding to environmental cues, such as changes in diet fat and temperature, at the transcriptional level (Luo et al. 2020; Shu et al. 2019). Atlantic salmon is an interesting natural model system for investigating the molecular mechanisms that directly link sexual maturation, energy acquisition, and environmental changes given that *vgll3* serves as a major activating transcription co-factor of the Hippo pathway, its tight linkage to maturation and adipogenesis processes (Ahi, Verta, et al. 2024b).

A growing body of evidence highlights the involvement of the Hippo pathway in various sex-dependent biological processes across vertebrates, such as size dimorphism in fish and reptiles (Wang et al. 2021; Zhu et al. 2022), innate immunity (Uppala et al. 2024; Kennicott & Liang 2024), adrenal gland development in mammals (Levasseur et al. 2019), and gonadal development and maintenance in fish (Cao et al. 2023). Of particular interest among the components of the Hippo pathway is *VGLL3*, which has been identified as a key regulator of sex-biased autoimmune responses in humans (Liang et al. 2017). Here, *VGLL3* influences a network of genes involved in various metabolic, developmental, and reproductive functions (Liang et al. 2017). This leads to hypotheses about its potential evolutionary significance in maintaining sex-specific metabolic homeostasis under metabolic stress, with implications for autoimmune-related pathological conditions (Pagenkopf & Liang 2020). However, the molecular details of such sex-dependent roles for *VGLL3* in regulating sexual maturation have not extensively been explored in any vertebrate species. Therefore, the *early* and *late* alleles of *vgll3* in Atlantic salmon present a unique opportunity to investigate this aspect in greater depth.

In this study, we employed a custom-made Nanostring gene expression panel to analyze the expression patterns of 333 genes, encompassing components of the Hippo pathway and their associated interacting partners, in the brain, ovary, and adipose tissue of immature female Atlantic salmon with homozygous *early* or *late vgll3* genotypes. We compared the patterns of gene expression in females to previously published data in respective male tissues collected at a seasonal stage coinciding with the onset of sexual maturation in males carrying the *vgll3*EE* genotype (Ahi, Verta, et al. 2024b). By doing so, we provide deeper insights into the *vgll3*-dependent molecular processes underlying its sex-dependent effects on sexual maturation.

## Materials and methods

### Fish material and tissue sampling

Individuals used in this study include eight males at the *immature-2* stage reported in Ahi et al. 2024a, b as well as eight females reared in the same tanks as these males. Details of the rearing and male sampling can be found in Verta et al., 2020 and Ahi et al. 2024b, respectively. Briefly, individuals from the same population (Oulujoki) and cohort used in Verta et al. (2020) provided access to individuals with known *vgll3* genotypes (see Verta et al. (2020) for details on crossing and rearing). Immature females reared in the same tanks were sampled at 1.5 years post-fertilization. Following euthanization by anesthetic overdose of MS222, various tissues, including visceral adipose tissue, brain, and ovary, were collected from the females during the summer (July 4-17). This time point was chosen as some males at this stage start to exhibit early signs of phenotypic maturation, while there were no phenotypic signs of maturation in any female (GSI of all females < 0.1). The female individuals had an average mass of 41.3 g (range 30.1-68.8 g) and an average length of 19.4 cm (range 15.0-20.3 cm) compared to the males with an average mass of 33.7 g (range21.1-76.6 g) and an average length of 17.3 cm (range 14.0-18.5 cm).

### RNA extraction and the Nanostring nCounter mRNA expression panel

In total, RNA was extracted from 24 tissue samples from eight immature females: eight samples each of visceral adipose tissue, brain, and ovary. RNA extraction was performed using a NucleoSpin RNA kit (Macherey-Nagel GmbH & Co. KG) as reported in Ahi et al. 2024b, and female samples were randomized among the male samples from Ahi et al. 2024a, b used in this study. RNA extraction followed the manufacturer’s instructions, including a built-in DNase step to remove residual gDNA. The extracted RNA from each sample was eluted in 50 µl (adipose and brain) and 80 µl (ovary) of nuclease-free water. RNA quantity was measured with a NanoDrop ND-1000 (Thermo Scientific, Wilmington, DE, USA), and quality was assessed with a 2100 BioAnalyzer system (Agilent Technologies, Santa Clara, CA, USA). The RNA integrity number (RIN) was > 7 for all samples. For each extraction, 100 ng of total RNA was used for the hybridization step in the Nanostring panel.

NanoString nCounter is a multiplex nucleic acid hybridization technology that enables assessment of RNA expression of several hundred genes simultaneously (Goytain & Ng 2020). As it requires only small RNA amounts with lower quality than RNA-seq, lacks an amplification step, and detects very low RNA expression levels, it is particularly attractive for ecological and evolutionary research (Kurko et al. 2020; Ahi, Verta, et al. 2024b; Richman et al. 2024; Ellison et al. 2021). The NanoString panel used here extends the panel used in Kurko et al. (2020) by adding more than 140 genes for a total of 337 genes. This panel includes an extensive list of Hippo pathway components and interacting partners (Kurko et al. 2020). The panel also included probes for age-at-maturity-associated genes in Atlantic salmon: *vgll3a* and *six6a* (on chromosome 25 and 9, respectively) and their paralogs *vgll3b* and *six6b* (on chromosome 21 and 1, respectively) as well as probes for other functionally relevant genes involved in metabolism, adipogenesis, and sexual maturation. Further details on gene/paralog selection and naming are available in Kurko et al. (2020), and gene accession numbers, symbols, full names, and functional categories as well as RNA hybridization procedures can be found in Ahi et al. (2024b).

### Data analysis

In the ovary, all nine candidate reference genes in the panel, including *actb*, *ef1aa*, *ef1ab*, *ef1ac*, *gapdh*, *hprt1*, *prabc2a*, *prabc2b* and *rps20*, were used for data normalization due to their low coefficient of variation (CV) values across the samples. In the adipose tissue, seven genes (excluding *actb* and *gapdh*) were chosen for normalization as they also exhibited low CV values. For the brain, the same set of reference genes was used, excluding *gapdh* because of its high expression variation across samples in this tissue. The raw count data from the Nanostring nCounter mRNA expression analysis was normalized using an RNA content normalization factor. This factor was calculated based on the geometric mean of selected reference gene counts for each tissue. Following normalization, a quality control check was conducted, and all samples met the default threshold as determined by the nSolver Analysis Software v4.0 (NanoString Technologies). During data analysis, the mean of the negative controls was subtracted, and normalization of the positive controls was performed using the geometric mean of all positive controls. A normalized count value of 20 was set as a background signal threshold. Below-average background signals were detected in 121, 82, and 77 genes across the samples in adipose, brain, and ovary, respectively, and were removed from further analyses. Sex-specific differential expression and Weighted Gene Coexpression Network Analysis (see below) were only investigated in the two tissues that are analogous between males and females (brain and adipose). Differential expression analysis was conducted using the log-linear and negative binomial model (lm.nb function) as implemented in Nanostring’s nSolver Advanced Analysis Module (nS/AAM). Sexes and genotypes were selected as predictor covariates in the model, as suggested by nS/AAM. Multiple hypothesis testing adjustment was performed using the Benjamini-Yekutieli method (Benjamini & Yekutieli 2001) within the software, and adjusted p-values < 0.05 were considered significant (Supplementary file 1).

The Weighted Gene Coexpression Network Analysis (WGCNA version 1.68) R-package (version 5.2.1) was implemented to identify gene co-expression modules (GCM) (Langfelder & Horvath 2008). Since our main interest was the comparison between sexes, all samples from both genotypes (for the adipose tissue and brain, separately) were used as biological replicates, providing sufficient statistical power for WGCNA. To identify sample relationships, hierarchical clustering of samples based on gene expression was conducted. Coexpression networks were constructed via seven steps described in (Ahi, Verta, et al. 2024b). Then, a conditional coexpression analysis was conducted as in Singh et al., 2021). Coexpression networks were constructed for each sex separately to identify the preservation of female modules in the male network and vice versa. A soft power of 8 was used to construct the adjacency matrix. Finally, module preservation statistics were calculated using WGCNA to test how the density and connectivity of modules defined in the reference dataset (e.g., female brain) were preserved in the query dataset (e.g., male brain) (Langfelder et al. 2011). A permutation test was implemented in the query network to calculate Z-scores and individual Z-scores from 200 permutations were summarized as a Z-summary statistic.

To further characterize the GCMs identified through WGCNA, we used WebGestalt (Elizarr Ar As et al. 2024) to examine similarities and differences between sexes in the biological processes associated with the genes within each module. Specifically, we tested for changes in module GO-association between the sexes as follows. We first identified GO-associations for each module using a gene-set overrepresentation test at a false discovery rate (FDR) of < 0.05, with a specific threshold for Gene Ontology/Biological Process (GO/BP) level 2 inclusion and all protein-coding genes as background. We then compared the GO/BP associations of each module between the sexes. This was performed for each tissue separately. To predict potential gene interactions and identify key genes with the highest number of interactions (interacting hubs), the identified differentially expressed genes in each comparison were converted to their conserved orthologs in humans (providing the highest amount of validated/studied interactome data in vertebrates) and used as input for STRING version 12.0 using the medium confidence level for predicting each interaction/molecular connection (Szklarczyk et al. 2023). The predicted interactions between genes were derived from data on structural similarities, cellular co-localization, biochemical interactions, and patterns of co-regulation.

## Results

### Female gene expression differences between *vgll3* genotypes

Assessment of expression differences between *vgll3* genotypes in females revealed 15 differentially expressed genes (DEGs) in the brain, 16 genes in the ovary and 29 genes in adipose tissue. In the brain, 10 out of 15 DEGs showed higher expression in *vgll3*EE* genotype individuals (Fig. 1A). A query of molecular interactions revealed three genes, *rhoaf, pparga and frmd6a*, showing direct interaction with *yap1* (specified with connecting lines between the genes in Fig. 1B). Among these three genes, *frmd6a* showed lower expression in *vgll3*EE* individuals whereas the two other genes (*rhoaf* and *pparga*) had higher expression in *vgll3*EE* individuals (Fig. 1B). Similarly in the ovary, 10 out of 16 DEGs showed higher expression in *vgll3*EE* genotype individuals (Fig. 1A). The predicted interactions revealed six genes (*arrb1*, *amotl2a*, *pax3b*, *snai2b*, *tead1a* and *tead3a*) with direct molecular interaction with *yap1*. In addition, three of these genes (*amotl2a*, *tead1a* and *tead3a*) had a direct interaction with *vgll3* (Fig. 1B). Three of the interacting genes, *arrb1*, *snai2b*, and *tead3a*, had higher expression in *vgll3*EE* individuals whereas the remaining three had lower expression in this genotype. Unlike the brain and ovary, all 29 DEGs identified in adipose tissue showed lower expression in *vgll3*EE* genotype individuals (Fig. 1A). The interaction query identified seven adipose DEGs (*ajubab*, *ets1e*, *foxo1c*, *lats2a*, *kdm5bc*, *tead3a* and *wwtr1b*) with direct interactions with *yap1*, whereas four genes (*akap11a, ets1e*, *tead3a* and *wwtr1b*) had a direct interaction with *vgll3* (Fig. 1B). One of these differentially expressed genes with direct connection with *yap1*, *foxo1c*, formed an interacting hub, and three of its interacting genes, *pparaa, ppargc* and *esrrab*, appeared to make further interacting hubs by connecting to other genes (Fig. 1B). We did not find any DEGs overlapping between the tissues (Fig. 1E), and adipose tissue was the only tissue where *vgll3a* was differentially expressed, with lower expression in *vgll3*EE* genotype individuals.

**Figure 1:**
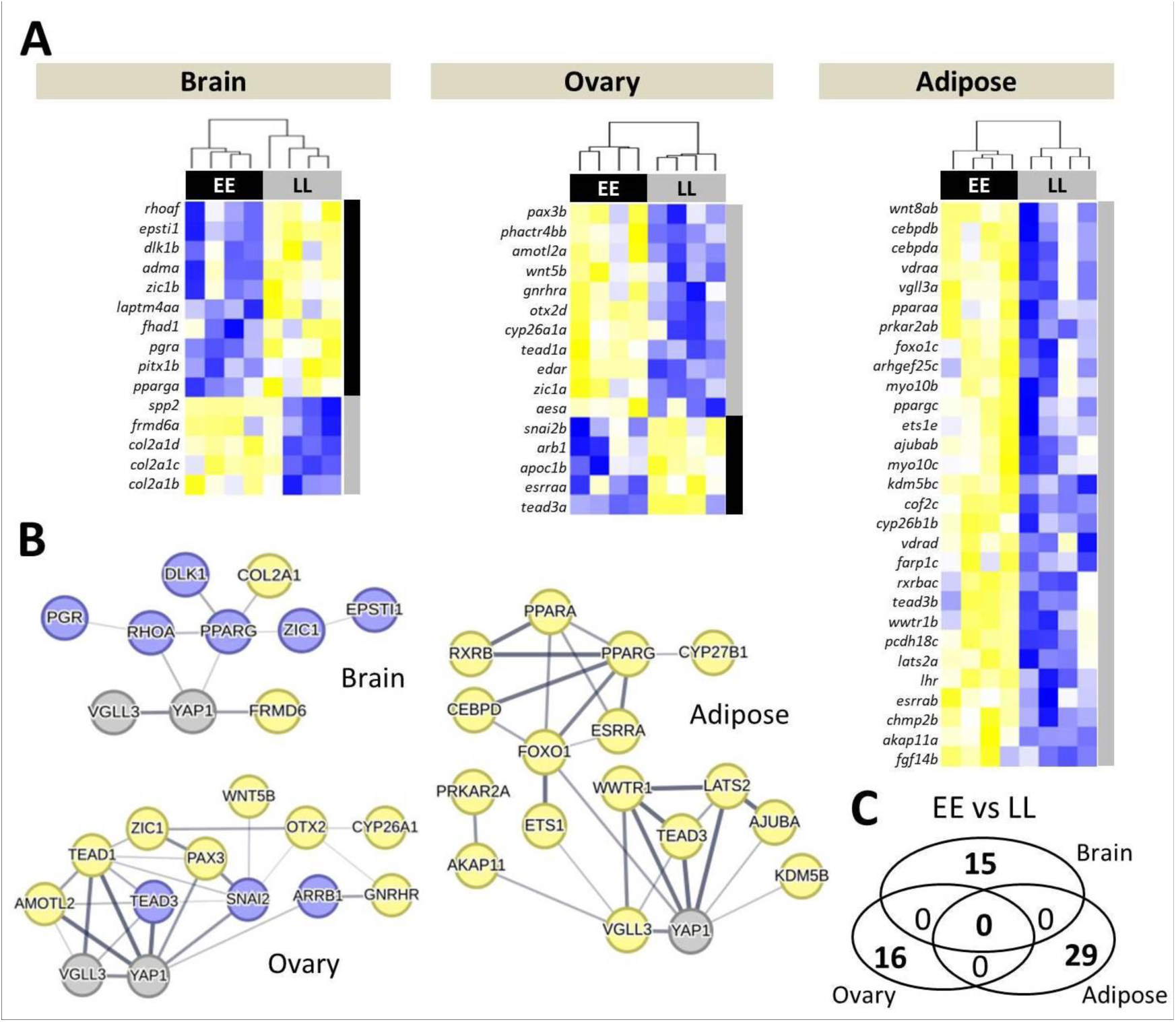
Differentially expressed genes in three tissues of female Atlantic salmon with alternative *vgll3* genotypes and their predicted interactions. Heatmaps represent differentially expressed genes between *vgll3* genotypes in three tissues (**A**) and their respective predicted interactions in each tissue using STRING v12 (http://string-db.org/) (**B**). The thickness of the connecting lines between the genes indicates the probability of the regulatory/functional interactions. Blue and yellow colors in A indicate higher and lower expression, and in B, higher and lower expression in *vgll3*EE* individuals, respectively. A Venn diagram showing the lack of of differentially expressed genes overlapping between the comparisons (**E**).

### Sex-specific gene expression differences in the brain

To identify gene expression differences between the female and male brain, we compared individuals from both sexes at a similar developmental time point in the summer (the *Immature 2* stage from Ahi et al. 2024), when some males, but no females, were starting to show phenotypic signs of maturation (see Methods). We identified 45 DEGs between female and male individuals when both *vgll3* genotypes were combined, and when considering *vgll3* genotypes separately, 23 and 33 DEGs were identified in female *vs* male *vgll3*LL* and *vgll3*EE* genotypes, respectively (Fig. 2A). Furthermore, we found a general tendency for DEGs to have higher expression in the brain of immature females whereby higher expression in females was observed in all of the 45 genes for the combined genotypes, 21 out of 23 DEGs for the *vgll3*LL,* and 29 out of 33 DEGs for the *vgll3*EE* comparisons (Fig. 2A). Across all three comparisons, expression patterns of three DEGs, *arhgef25b, cadm2a* and *frm6a*, were independent of *vgll3* genotype, and all these genes had a higher expression level in immature females compared to males (Fig. 2A). These results indicate higher transcriptional activity of the studied genes in the brain of immature females compared to males at this time of the life-cycle.

**Figure 2:**
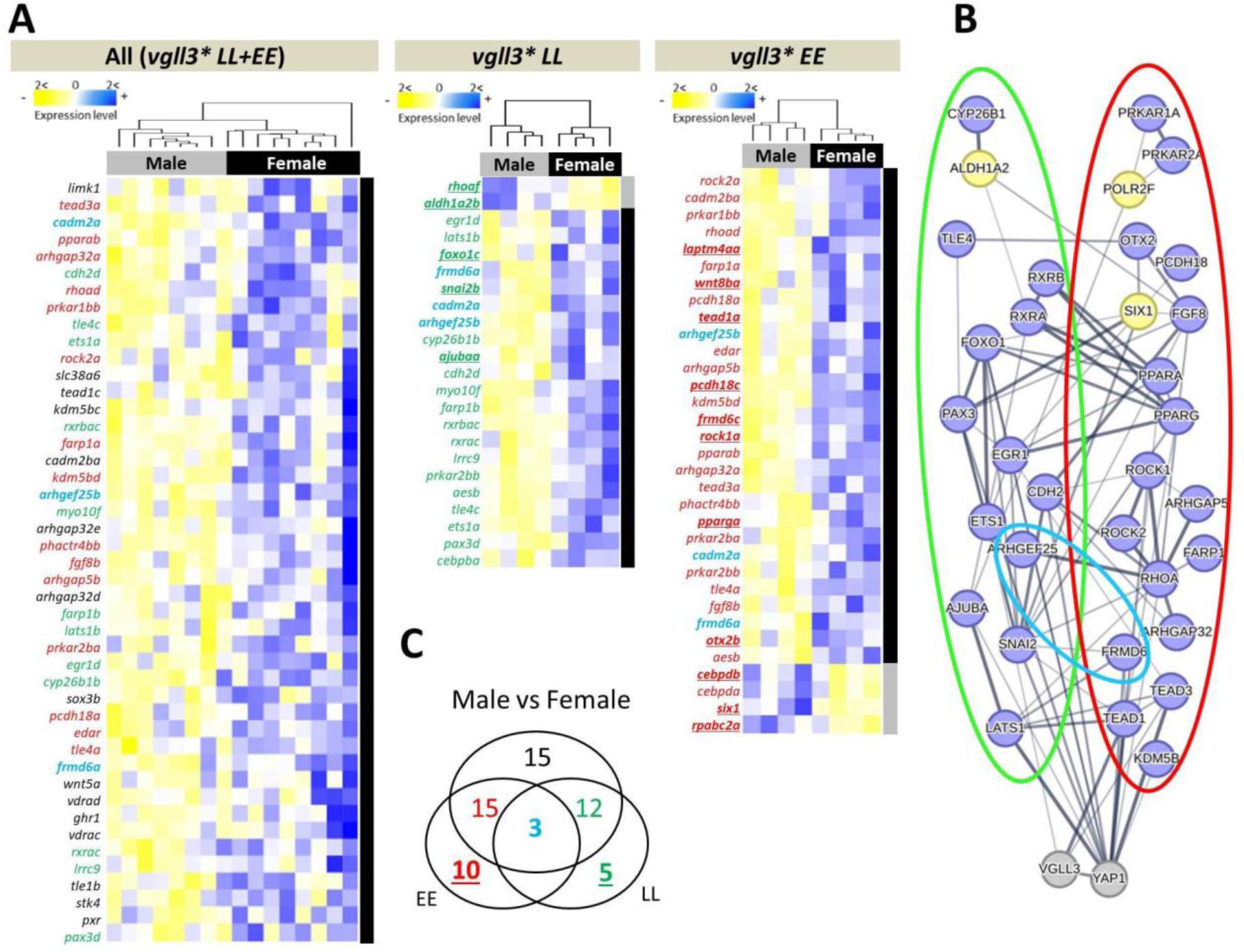
Differentially expressed genes between female and male Atlantic salmon and their predicted interactions in the brain. Heatmaps representing differentially expressed genes between the sexes in immature individuals with alternative *vgll3* genotypes pooled (**A**), and within *vgll3*LL* (**B**) and *vgll3*EE* genotypes (**C**). Predicted interactions between the overlapping genes, with green, red and blue rings indicating the genes in the Venn diagram (**D**). The thickness of the connecting lines between the genes indicates the probability of the interaction. Blue and yellow colors indicate higher and lower expression in female individuals, respectively. A Venn diagram showing the numbers of differentially expressed genes overlapping between the comparisons (**E**). The color coding of the numbers corresponds to the gene colors shown in the list of genes within the heatmaps.

We further investigated potential functional/molecular interactions between DEGs showing *vgll3* genotype-specific differential expression (colored numbers and circles in Fig. 2B-C). The predicted interactions between these genes revealed extensive and complex regulatory connections between the DEGs in each genotype (Fig. 2B). From the *vgll3* genotype-specific comparisons, 13 and 19 DEGs within *vgll3*LL* and *vgll3*EE* individuals, respectively, were found to be connected in the interaction network (circled in red and green in Fig. 2D). Moreover, except for *aldh1a2* within the *vgll3*LL* network (green circled), and *six1* and *rpabc2a/POLR2F* within the *vgll3*EE* network (red circled), all other genes in both networks had higher expression in females (Fig. 2B). Further, some of the genes in each genotype network had high numbers of interactions in the networks including *ets1a*, *egr1d*, *foxo1c*, *lats1b* and *snai2b* in the *vgll3*LL* interaction network, and *fgf8b*, *rhoad*, *tead1a* and *tead3a* in the *vgll3*EE* network (Fig. 2B). These findings imply that specific components of the Hippo pathway might be responsible for the elevated transcriptional activity observed in the brains of females compared to males.

### Sex-specific gene expression differences in adipose tissue

Similar comparisons to those conducted in brain tissue were conducted in adipose and seven DEGs were identified between female and male individuals when both *vgll3* genotypes were pooled. When considering *vgll3* genotypes separately, we found six DEGs in the *vgll3*LL* genotype and 23 DEGs in the *vgll3*EE* genotype individuals (Fig. 3A). Furthermore, we found that all DEGs in the pooled genotypes and in *vgll3*LL* had increased expression in females, whereas most DEGs in *vgll3*EE* (17 out of 23) had higher expression in males. No gene was differentially expressed in all comparisons (Fig. 3B). These results indicate significantly more pronounced and stronger sex-specific expression in the adipose tissue of individuals with the *vgll3*EE* genotype.

**Figure 3:**
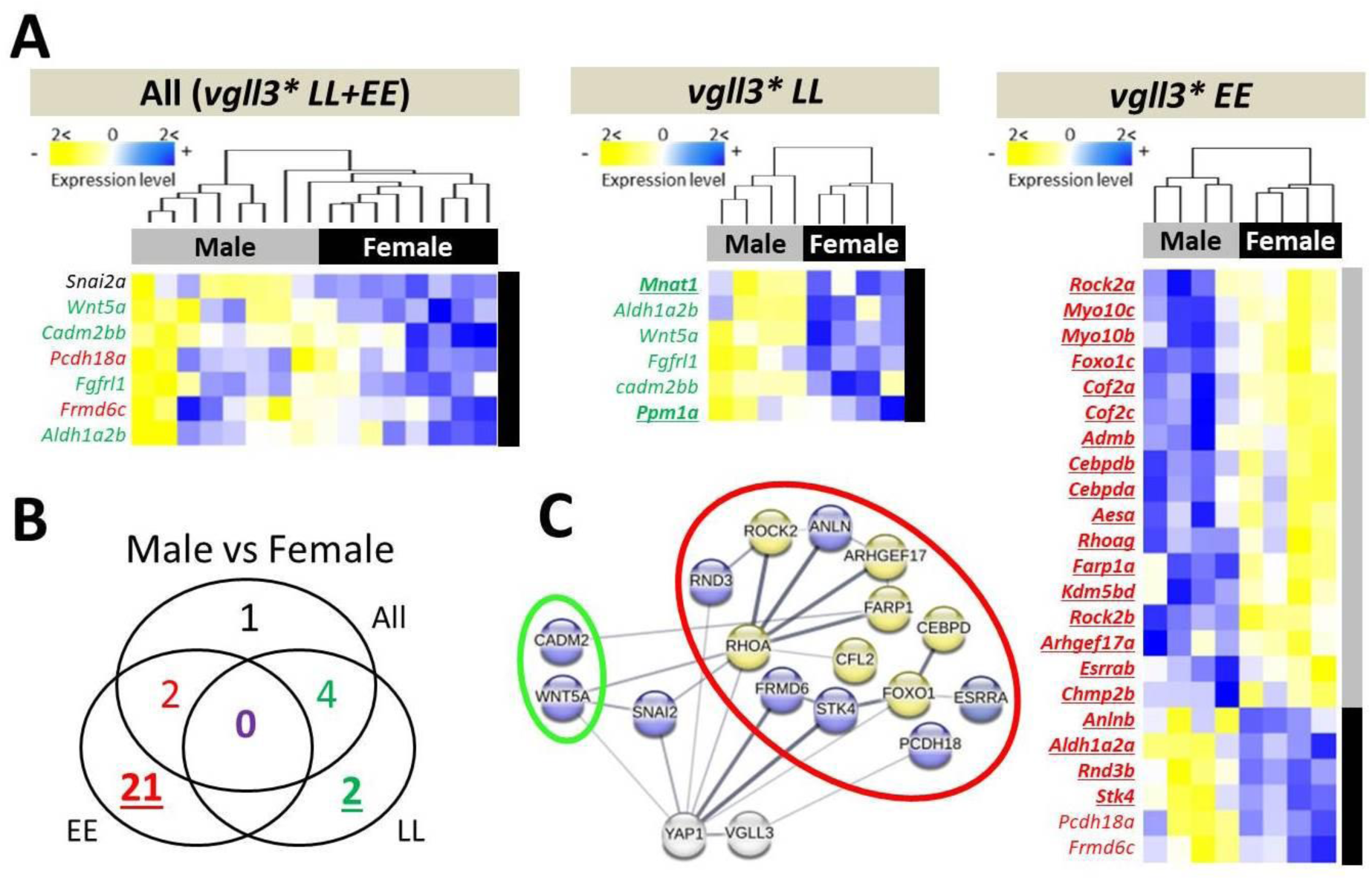
Differentially expressed genes between female and male Atlantic salmon and their predicted interactions in adipose tissue. Heatmaps representing differentially expressed genes between the sexes in the immature individuals across alternative *vgll3* homozygotes (**A**), and within *vgll3*LL* (**B**) and *vgll3*EE* genotypes (**C**). A Venn diagram showing the numbers of differentially expressed genes overlapping between the comparisons and the color coding of the numbers corresponds to the gene colors shown in the list of genes within the heatmaps (**D**). Predicted interactions between the overlapping genes, with green and red rings indicating the genes in the Venn diagram. The thickness of the connecting lines between the genes indicates the probability of the interaction (**E**). Genes colored in blue and yellow in A indicate higher and lower expression, and in C, higher and lower expression in females, respectively.

We further investigated potential functional/molecular interactions between DEGs showing *vgll3* genotype-specific differential expression (colored numbers in Fig. 3B). In the *vgll3* genotype-specific comparisons, two and 13 DEGs within *vgll3*LL* and *vgll3*EE* individuals, respectively, were found to be connected in the interaction network (circled in red and green in Fig. 2C). While both genes from the *vgll3*LL* comparison, *cadm2bb* and *wnt5a*, had higher expression in females compared to males, seven genes from the *vgll3*EE* comparison showed lower expression in the females (Fig. 2C). A few genes in each genotype-specific sex comparison had a high number of interactions in the networks, including *wnt5a* in the *vgll3*LL* network, as well as *rhoag*, *foxo1c*, and *stk4* in the *vgll3*EE* network (Fig. 2C). Among the genes with a high number of interactions in the predicted network, *rhoag* and *foxo1c* showed lower expression in females. Importantly, two key components of the Hippo pathway, *frmd6* and *stk4/mst1*, had the strongest predicted interaction with *yap1* and both showed increased expression in females with the *vgll3*EE* genotype. Taken together, these findings suggest that the pronounced transcriptional differences observed in the adipose tissue of *vgll3*EE* individuals are likely mediated by key components of the Hippo pathway, such as *frmd6* and *stk4/mst1*.

### Identification of sex-specific gene coexpression modules in the brain

In order to gain a better overview of sex-specific transcriptional differences of Hippo pathway components and their known interacting genes in the brain, we applied network-based co-expression analyses in which changes between the sexes in each network could be tracked. To do this, we first built gene coexpression modules (GCMs) in the brain of each sex and then investigated the preservation of the identified GCMs between the sexes. In other words, we defined the GCM in one sex and then assessed the preservation of its modules in the other sex. We identified five brain GCMs for females (Fig. 4A and B) of which one GCM (brown) showed relatively high preservation (Zsummary > 2) in males, i.e. most of the genes in this GCM have significant expression correlations in both sexes. Three of the GCMs (yellow, green and blue) showed a low to moderate level of preservation between the sexes (Zsummary = 0 - 2); and the turquoise GCM, containing the highest number of genes (63 genes), showed the lowest level of preservation (Fig. 4B). Next, a gene set enrichment analysis was conducted for the GCM with the lowest level of preservation between females and males (turquoise) in order to provide insights into the biological processes associated with male vs. female differences. The most common biological processes of the genes in the least preserved (turquoise) GCM included regulation of vitamin D biosynthesis, cell-cell junction assembly, Hippo signaling pathway, cellular response to lipid and steroid hormone mediated signaling pathway (Fig. 4C). Almost half its genes in this turquoise GCM showed no coexpression preservation in males (genes lacking color in Fig. 4C). Removal of unpreserved genes in the turquoise GCM led to loss of significance of three GO terms; Hippo signaling pathway, cellular response to lipid and steroid hormone mediated signaling pathway (non-colored GOs in Fig. 4C). Knowledge-based interactome prediction using genes within the turquoise GCM was performed in order to identify potential interactions between the genes as well as hub genes with highest number of interactions. The prediction of interactions between the genes within this GCM revealed that several genes among those lacking coexpression preservation directly interacted with *vgll3/yap1* (represented with lines directly connecting the non-colored genes with *vgll3/yap1*; Fig. 4D). These genes include *ets1*, *tead1*, *tead3a* and *wwtr1b* showing direct interactions with *vgll3* as well as *egr1*, *kdm5b*, *pax3*, *rhoa*, *rock1*, *snai2b*, *snai1*, *stk3*, *tead1*, *tead3a* and *wwtr1b* showing direct interactions with *yap1* (Fig. 4D).

**Figure 4.**
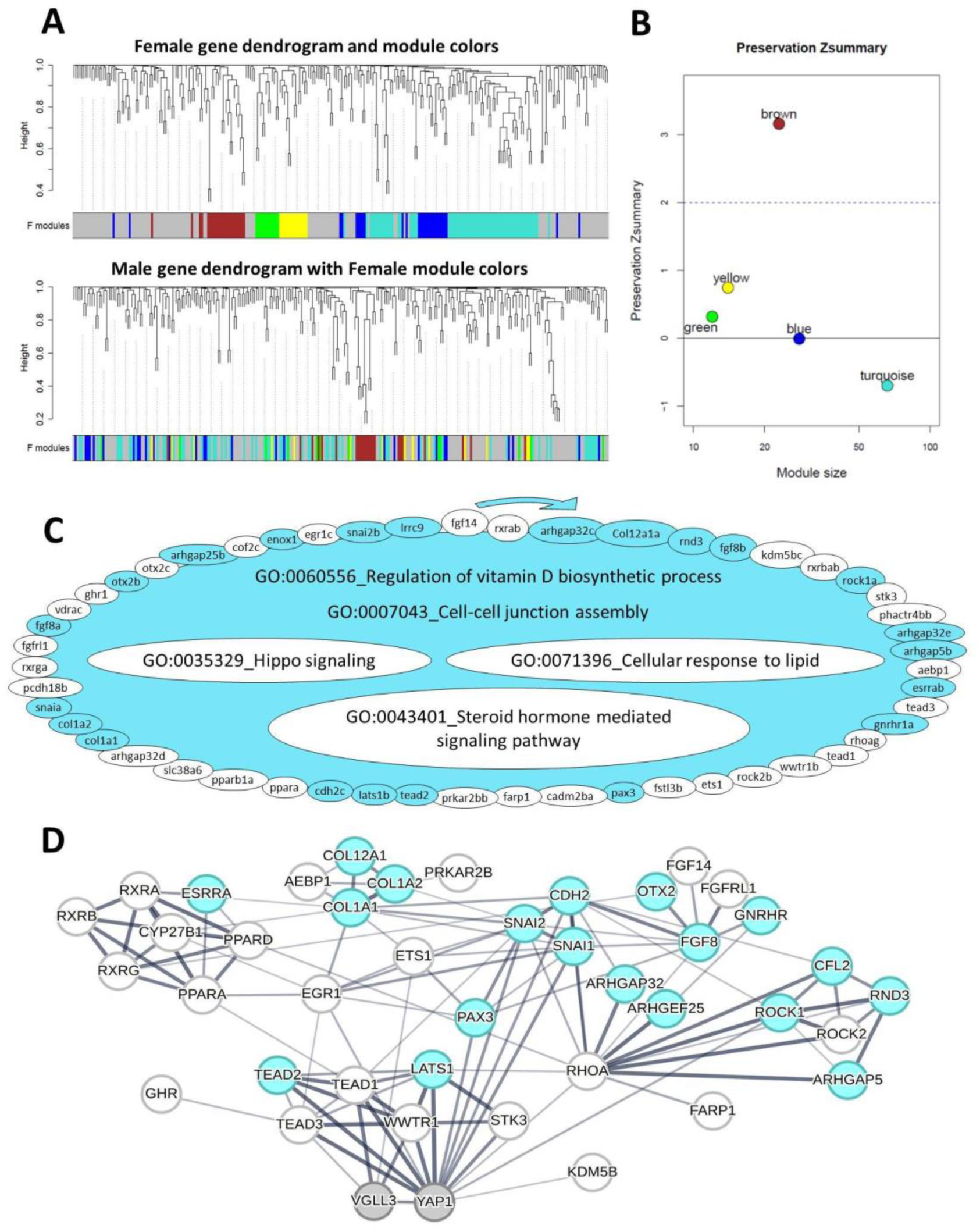
Coexpression modules in the brain of female Atlantic salmon. Visual representation of female module preservation in male individuals. The dendrograms represent average linkage clustering tree based on topological overlap distance in gene expression profiles. The lower panels of the dendrograms represent colors that correspond to the female clustered coexpression modules (GCMs). Top: female GCMs with assigned colors. Bottom: visual representation of the lack of preservation of female GCMs genes in male individuals (**A**). Preservation Zsummary scores in the male GCMs for female GCMs (colors represent female GCMs). Zsummary < 0 represents lack of preservation (dotted blue line) and Zsummary 0 - 2 implies moderate preservation (**B**). The genes in turquoise GCM identified in female brain with least preservation in males. The genes without color in the module are those showing no preserved expression correlation in male individuals and the clockwise arrows above the GCM indicate the direction of genes with highest to lowest expression correlations with other genes within the GCM. In the turquoise GCM, the top enriched GOs are represented, and GOs without color were no longer enriched after removal of the genes without colors (**C**). Predicted interactions between the genes within the turquoise GCM. Increasing thickness level in the connecting lines between the genes indicates a higher probability of the interaction (**D**).

We found four GCMs in males and among them, the red GCM showed the lowest level of preservation (Zsummary < 0) compared to females (Fig. 5A and B). The red GCM was also the largest with 42 co-expressed genes and more than half of these genes showed no coexpression preservation in females compared to males (genes lacking color in this GCM in Fig. 5C), suggesting this GCM to be of most interest for identifying genes important for male vs female differences. The gene set comparison of the least preserved red GCM identified four GOs including regulation of tight junction assembly, Hippo signaling pathway, developmental growth and hormone-mediated signaling pathway (Fig. 5C), suggesting these processes to be important in determining male vs. female differences. Furthermore, removal of unpreserved genes in the red GCM led to absence of significance of all these GOs (non-colored GOs in Fig. 5C).The prediction of interactions between the genes within the red GCM revealed that six genes that lost their coexpression preservation, *frmd6*, *rhoa*, *rock1*, *sav1*, *snai* and *tead3*, had direct interaction with *yap1* whereas two unpreserved genes, *ets1* and *tead3* had direct interactions with *vgll3* (Fig. 5D). Importantly, *yap1* itself lost coexpression preservation in the red GCM, indicating the involvement of *yap1*, the major inhibitor of the Hippo pathway, in sex differences in the lowest preserved GCM. This suggests that the observed male-specific transcriptional pattern in the brain might be mediated by *yap1*-dependent differential activity of the Hippo pathway.

**Figure 5.**
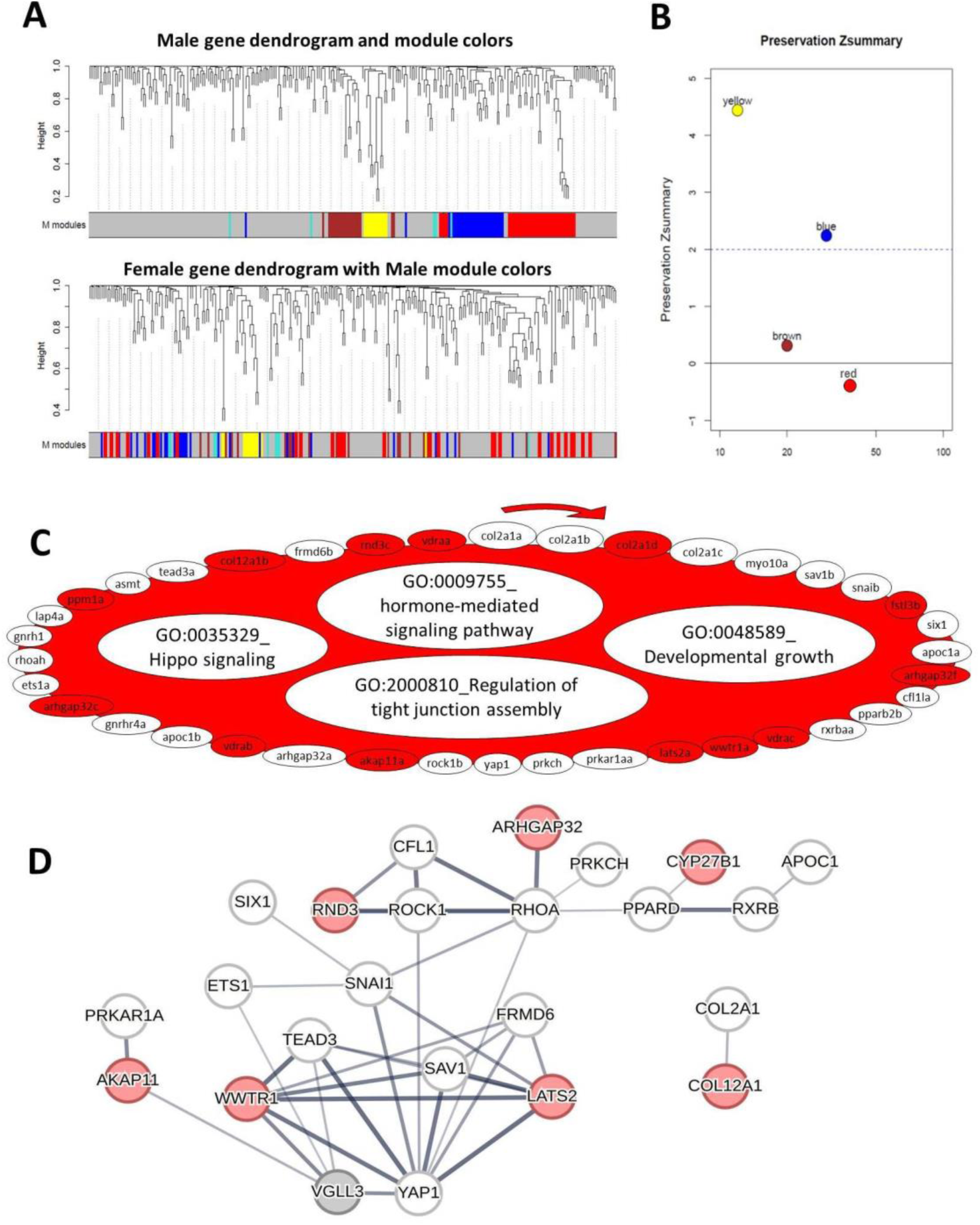
Coexpression modules in the brain of male Atlantic salmon. Visual representation of male module preservation in female individuals. The dendrograms represent average linkage clustering tree based on topological overlap distance in gene expression profiles. The lower panels of the dendrograms represent colors that correspond to the male clustered coexpression modules (GCMs). Top: male GCMs with assigned colors. Bottom: visual representation of the lack of preservation of male GCMs genes in female individuals (**A**). Preservation Zsummary scores in the female GCMs for male GCMs (colors represent male GCMs). Zsummary < 0 represents lack of preservation (dotted blue line) and Zsummary between 2 and 6 implies moderate preservation (**B**). The genes in the red GCM identified in males with least preservation in females. The genes without color in the module are those showing no preserved expression correlation in female individuals and the clockwise arrows above the GCM indicate the direction of genes with highest to lowest expression correlations with other genes within the GCM. In the red GCM, the top over-represented GOs are listed, and GOs without color were no longer enriched after removal of the genes without colors (**C**). Predicted interactions between the genes within the red GCM. Increasing thickness level in the connecting lines between the genes indicates a higher probability of the interaction (**D**).

### Identification of sex-specific gene coexpression modules in the adipose tissue

Gene coexpression module analyses were conducted for the adipose tissues of both sexes as described for the brain (see above). After building the GCMs for the adipose tissue of each sex, we investigated the preservation of the identified GCMs between the sexes and found no unpreserved GCM in male adipose tissue, indicating that the identified GCMs in males occur in both sexes. However, among the seven GCMs identified in the females (Fig. 6A and B), one GCM (yellow) showed a very low level of preservation (Zsummary < 0) in males and it was thus investigated further using gene-set enrichment as above. We detected two over-represented GO terms, namely regulation of cAMP-dependent protein kinase activity and Hippo signaling pathway (Fig. 6C). In the yellow GCM, we also found that almost half of the genes showed no coexpression preservation in males (nine out of 19 genes lacking color in this GCM in Fig. 6C). Removal of unpreserved genes in the yellow GCM led to loss of significance of the GO term associated with the Hippo signaling pathway (the non-colored GO in Fig. 6C). Knowledge-based interactome prediction using genes within the yellow GCM revealed several hub genes (e.g., *amotl2*, *lats1*, *rhoa*, *fgf2* and *wnt5a*) (Fig. 6D). Furthermore, among those genes not showing coexpression preservation, one gene, *kdm5b*, had direct predicted interaction with *vgll3* and three genes, *kdm5b*, *lats1* and *fgf2*, had direct interactions with *yap1* (represented with lines directly connecting the non-colored genes with *vgll3/yap1*; Fig. 6D). Among the genes directly interacting with *vgll3* and/or *yap1*, three genes; *sav1*, *amotl2*, and *lats1*; show strong interactions and acting as upstream regulators of the Hippo pathway (all are upstream inhibitors of *yap1*).

**Figure 6.**
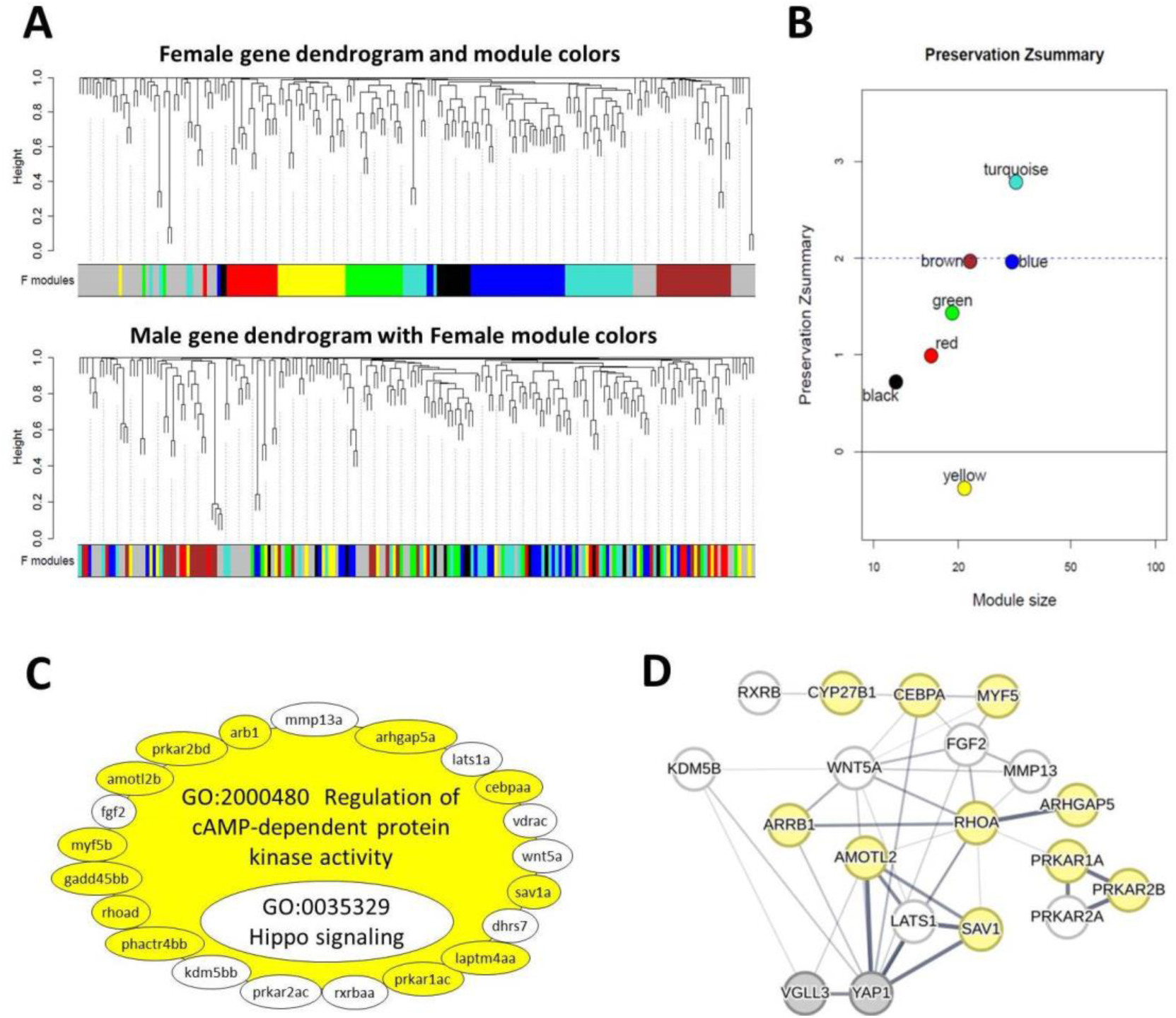
Coexpression modules in female Atlantic salmon adipose tissue. Visual representation of female module preservation in male individuals. The dendrograms represent average linkage clustering tree based on topological overlap distance in gene expression profiles. The lower panels of the dendrograms represent colors that correspond to the female clustered coexpression modules (GCMs). Top: female GCMs with assigned colors. Bottom: visual representation of the lack of preservation of female GCMs genes in male individuals (**A**). Preservation Zsummary scores in the male GCMs for female GCMs (colors represent female GCMs). Zsummary < 0 represents lack of preservation (dotted blue line) and Zsummary 0 - 2 implies moderate preservation (**B**). The genes in yellow GCM identified in female adipose tissue with least preservation in males. The genes without color in the module are those showing no preserved expression correlation in male individuals and the clockwise arrows above the GCM indicate the direction of genes with highest to lowest expression correlations with other genes within the GCM. In the yellow GCM, the top enriched GOs are represented, and GOs without color were no longer enriched after removal of the genes without colors (**C**). Predicted interactions between the genes within the yellow GCM. Increasing thickness level in the connecting lines between the genes indicates a higher probability of the interaction (**D**).

Since only *lats1* does not show co-expression preservation, this suggests that *lats1* may play a critical role in mediating the regulatory effects seen in this tissue. Taken together, these results suggest that the overall sex-specific differences in the expression of Hippo pathway components and its interacting partners might be influenced by *lats1* transcription in the adipose tissue.

## Discussion

In this study, we explored how [sex and] the genotype of a major maturation age gene, *vgll3*, interplay[s] with tissue in driving sex-specific expression patterns potentially associated with the onset of maturity in Atlantic salmon. We did this by profiling the expression of known components of the Hippo pathway and their interacting partners in brain, adipose, and gonad tissues at a stage when some males begin to exhibit signs of pubertal initiation while other males and all females remain immature. In general, we observed more extensive *vgll3* genotype effects on gene expression in adipose tissue of females compared to the brain and ovary. Compared to the other two tissues, expression differences in adipose tissue seemed to be more linked with Hippo pathway signaling, as several components of this pathway, including *vgll3a* itself, were differentially expressed between alternative *vgll3* genotypes. These results are concordant with earlier research in mammals suggesting the Hippo pathway plays a pivotal role in balancing adipocyte proliferation vs. differentiation (Ardestani et al. 2018). For example, *vgll3* has been reported as an inhibitor of adipocyte differentiation in mice (Halperin et al. 2013), and also the activity of Yap has been suggested to be indispensable during adipogenesis (Ardestani et al. 2018). Additionally, a major Hippo pathway kinase, encoded by *lats2*, which was differentially expressed in female adipose tissue in our study, is known to promote the lipolysis process in mouse adipocytes (El-Merahbi et al. 2020). Our recent findings in male Atlantic salmon also imply that *vgll3* and its associated Hippo pathway have extensive effects on transcriptional changes in adipose tissue in relation to sexual maturation, as well as linking adipogenesis and seasonal changes in this species (Ahi, Verta, et al. 2024b).

In contrast to adipose tissue, in the female brain, there were fewer DEGs, and none of the major components of the Hippo pathway were found to be differentially expressed between the *vgll3* genotypes. This suggests a potentially significant difference in the functional role of Hippo pathway signaling between the brain and adipose tissue in females, at least at this developmental time point. In contrast, at the same immature stage, the brain of males showed very distinct transcriptional activation of the Hippo pathway between the genotypes (Ahi, Verta, et al. 2024a). In males, *tead2*, encoding a major transcription factor of the Hippo pathway, and three interacting partner genes of the Hippo pathway (*kdm5b/jarid1b*, *mc4ra*, and *foxo1c*) that play roles in the central regulation of the onset of puberty (Farooqi et al. 2003; Lomniczi et al. 2015; Liu et al. 2022), had higher expression in the brain of individuals with the *vgll3*EE* genotype (Ahi, Verta, et al. 2024a). None of these genes were differentially expressed in the female brain between the *vgll3* genotypes, indicating a sex-specific difference in the involvement of Hippo pathway components in central sexual maturation signals at this developmental timepoint (Fig. 1). However, four additional genes—*dlk1b*, *pgra*, *rhoaf*, and *zic1b*—which encode other interacting partners of the Hippo pathway, were found to have higher expression in the brains of females with the *vgll3*EE* genotype. This is particularly striking because the orthologs of these genes (*DLK1*, *PGR*, *RHOA*, and *ZIC1*) are all known to be involved in the central regulation of pubertal onset in human (Cassin et al. 2023; Swift-Gallant et al. 2015; Dauber et al. 2017; Sklirou & Lahr 2023). Furthermore, two of these genes, *DLK1* and *PGR*, have also been demonstrated to have sex-specific roles during puberty.

In the ovary, we found a lower number of DEGs between *vgll3* genotypes than in adipose tissue. However, unlike the brain, major components of the Hippo pathway were differentially expressed, such as *amotl2a*, *arrb1*, *tead1a*, and *tead3a*. Recent studies have shown that the core components of the Hippo pathway play important roles in mammalian ovarian physiology, including ovarian development, follicle development, and oocyte maturation (reviewed by Clark et al. 2022). For instance, the higher expression of *arrb1* in the ovary of fish individuals with the *vgll3*EE* genotype could thus indicate enhanced ovarian development, as *arrb1* is involved in cellular responses to hormones and growth factors and is an important marker of developing ovary (Zhou et al. 2019). Among the interacting partners of the Hippo pathway, *snai2b*, a known marker of primordial ovarian follicles (Amoushahi & Lykke-Hartmann 2021), also showed higher expression in the ovary of *vgll3*EE* individuals. Another notable gene with increased expression in the ovary of *vgll3*EE* individuals was *esrra* (*nr3b1*), which encodes an orphan estrogen receptor with an important role in angiogenesis during ovarian development (Guzmán et al. 2021). Moreover, we found reduced expression of *cyp26a1*, which encodes an enzyme that inhibits ovarian development by blocking retinoic acid (RA) signaling, in the ovary of individuals with the *vgll3*EE* genotype (Demczuk et al. 2016). These findings suggest potential differences in Hippo pathway-mediated ovarian development between the *vgll3* genotypes in Atlantic salmon, with possibly more advanced ovarian development in *vgll3*EE* individuals. However, future studies on ovarian development, including time series analyses from immature to mature females, similar to the recent work focused solely on *vgll3* expression (Åsheim et al. 2024), are necessary to fully understand these differences. Such studies should explore not only the expression of the entire components of the pathway but also the changes at cellular-level, offering deeper insights into the regulatory interactions that govern ovarian maturation.

### Gene expression differences suggest differences in lipolysis capacity and adipogenesis of females with distinct *vgll3* genotypes

We found lower expression of *vgll3a* in the adipose tissue of *vgll3*EE* females, consistent with previous findings in males where we observed differences in lipid content and gene expression patterns in liver and adipose tissues between the *vgll3* genotypes, suggesting *vgll3*EE* males may store larger lipid droplets in the spring/summer (many months before spawning time), whereas *vgll3*LL* individuals store in the autumn (House et al. 2025; Ahi, Verta, et al. 2024b). This supports the scenario whereby *vgll3*EE* individuals have a higher adipogenesis capacity in the spring in both sexes. However, a closer examination of the DEGs between the genotypes adds further details to the interpretation of the results. For example, *vgll3* and *TAZ* (*WWTR1*), both of which had reduced expression in *vgll3*EE* individuals, which are described as inhibitors of the terminal stage of adipocyte differentiation (Halperin et al. 2013; Jung et al. 2009). On the other hand, we also found lower expression of initial-stage adipogenesis markers, such as *ppargc*, *cebpda*, *ajubab* and *esrra*, in *vgll3*EE* individuals (Ijichi et al. 2007; Yan et al. 2022; Kusuyama et al. 2017; Chawla et al. 1994). Specifically, while pre-adipocyte differentiation might be at a lower level in *vgll3*EE* individuals (due to reduced *ppargc*, *cebpda*, *ajubab* and *esrra* expression), the terminal stage of adipocyte maturation might be promoted (due to reduced *vgll3a* and *taz/wwtr1b* expression) and the opposite may be true for *vgll3*LL* individuals. This complex transcriptional signature suggests that the genotype difference may lie in specific stages of adipogenesis (e.g., heterochrony) rather than in overall adipogenesis capacity. In other words in *vgll3*EE* individuals, the pre-adipocyte differentiation phase has already been completed, and the adipocytes have entered the terminal stage of differentiation. In contrast, in *vgll3*LL* individuals, the pre-adipocytes are still in the early stages of differentiation and have not yet progressed to their final stage. In addition, we found reduced expression of lipolysis factors, including *lats2*, *foxo1c*, and *pparaa*, in *vgll3*EE* female adipose (Chakrabarti & Kandror 2009; El-Merahbi et al. 2020; Guzmán et al. 2004). This reduction may indicate an increased lipid-storing capacity in the *vgll3*EE* genotype, as the expression of lipolysis markers is typically reduced when lipid storage is prioritized. Together, these findings suggest reduced pre-adipocyte differentiation and lipolysis capacity, alongside enhanced adipocyte maturation and lipid storage capacity in *vgll3*EE* females during the summer. This is concordant to findings in males at the same developmental time point (House, Debes, Holopainen, et al. 2023; Ahi, Verta, et al. 2024b).

### Extensive sex-specific differences in transcription of the Hippo pathway components in the adipose tissue of *vgll3*EE* individuals

Assessment of adipose tissue transcriptional differences between the sexes revealed more pronounced sex differences in *vgll3*EE* individuals compared to *vgll3*LL* individuals (Fig. 3). Sex-specific expression patterns in *vgll3*EE* individuals were observed in components of the Hippo pathway (such as *stk4*, *frmd6c*, and *pcdh18a*) as well as interacting partners of this pathway that play important roles in adipogenesis and lipolysis (such as *esrra*, *foxo1c*, *rhoag*, and *cebpda/b*). Among the Hippo pathway components, *stk4* is known to have an important role in adipogenesis, with increased activity leading to augmented adipose mass and obesity while reducing the energy expenditure of adipose tissue by impairing mitochondrial function (Cho et al. 2021). Thus, the higher expression of *stk4* in *vgll3*EE* females may indicate weaker energy expenditure performance compared to males with the same genotype, resulting in more adipose mass gain. Consistently, the lower expression of a *RhoA* paralog gene (*rhoag*) in *vgll3*EE* females indicates a higher capacity for adipogenesis and lipid droplet storage, as RhoA is a major suppressor of both processes in mammalian cells (Meyers et al. 2005). Moreover, a pre-adipocyte differentiation marker, *esrra* (Ijichi et al. 2007), was found to be induced in *vgll3*EE* females, concordant with increased adipogenesis in these females. Higher expression of a *RND3* paralog gene (*rnd3b*), encoding a key inhibitor of lipolysis (Dankel et al. 2019), and reduced expression of *foxo1c*, an inducer of lipolysis (Chakrabarti & Kandror 2009), in *vgll3*EE* females suggests potentially reduced lipolysis in their adipose tissue. These results suggest higher fat accumulation and adipogenesis in *vgll3*EE* females compared to the males with the same genotype, but potentially with a lower capacity for energy expenditure.

### Sex-specific links between the Hippo pathway and cAMP-dependent protein kinase activity in adipose tissue

Our co-expression analysis revealed that only one GCM identified in the female adipose tissue was not preserved in the males (yellow module in Fig. 6). This means that most of the gene expression correlations within the yellow module were absent in males, meaning the functional relationships between these genes were not maintained in the adipose tissue of males. This GCM in females included correlated expression of several components of the Hippo pathway (such as *arrb1*, *lats1b*, *amotl2a*, and *sav1*) and genes involved in the regulation of cAMP-dependent protein kinase A (PKA) activity (such as *prkar1a*, *prkar2a*, *prkar2b*, *cebpa*, *myf5*, *fgf2*, and *rhoa*). PKA activity is a major signal controlling lipid metabolism, particularly lipolysis (Carmen & Víctor 2006). For instance, in mice, the loss of *prkar1a* enhances lipolysis in adipose tissue and leads to rapid weight loss (Gangoda et al. 2014), while *prkar2b* function is required for adipocyte differentiation (Peverelli et al. 2013). In males, the correlation between the Hippo and PKA signals appears to be absent (Fig. 6C). The major Hippo component affected was *lats1b* (another inducer of lipolysis (Shen et al. 2022)), as it did not show expression correlation with the PKA components in the adipose tissue of males. Interestingly, the crosstalk between the Hippo and PKA signals is known to be mediated directly through the phosphorylation of LATS1 or indirectly through the activation of RhoA by PKA in mammals (Yu et al. 2013). These interactions can lead to synergistic activation of both signals in various tissues. Our result here suggests that the sex-specific mechanism predicted earlier (see above), by which females exhibit higher adipogenesis and lower levels of energy expenditure compared to males, might originate from this link between the PKA and Hippo signals that exists only in the female adipose tissue. This may also indicate that while females might store a larger amount of lipid and gain more fat mass, they may delay in utilizing these reserves, leading to later maturation overall.

### Gene expression differences suggest higher activity of the Hippo pathway in the brain of females compared to males

We found genotype-independent induced expression of *frmd6a*, encoding a major upstream regulator of the Hippo pathway, in the brain of immature females when compared to immature males (Fig. 2). *FRMD6/Willin* expression is tightly co-localized with *GHRH* (growth hormone-releasing hormone) in nerve cells, particularly in the nerves densely populated in the hypothalamus and anterior pituitary of vertebrates (Beck & Kressel 2020). GHRH is one of the earliest discovered hypothalamic factors involved in the sexually dimorphic pubertal timing of mammals (Argente et al. 1991). However, the regulatory connection between FRMD6 and GHRH during sexual maturation remains to be elucidated. FRMD6 is considered a potent inhibitor of YAP1 and an activator of the Hippo pathway (Angus et al. 2011), suggesting that the Hippo pathway is generally more active in the female brain than in males at this time point. Despite distinct transcriptional signatures between the *vgll3* genotypes, both genotypes reflected higher Hippo pathway activity in the female brain (e.g., increased expression of *lats1b* in females with the *vgll3*LL* genotype, and *tead1* and *tead3a* in females with the *vgll3*EE* genotype) (Fig. 2). Another noteworthy gene with genotype-independent induced expression in females was *cadm2a*, which encodes another conserved cell adhesion protein highly expressed during vertebrate brain development (Hunter et al. 2011). *CADM2* is also known as a factor linking psychological/behavioral traits and obesity, as well as the brain and adipose tissues (Morris et al. 2019). *CADM2* has been indicated to have sex-dependent expression during brain development and function in humans (Wingo et al. 2023). Finally, the third gene identified with genotype-independent induced expression in the female brain was *arhgef25b* (known as *GEFT* in mammals), encoding a Rho-GTPase enzyme required for neurite outgrowth, which is responsible for neuronal patterning and connections (Bryan et al. 2004). A study in clownfish found *arhgef25* to be one of the few sex-dependent genes during sexual transition, required during the female stage in this species (Casas et al. 2016). These findings suggest a potential molecular axis whereby the brain senses varying energy storage status in adipose tissue and responds in a sex-specific manner. This axis may be triggered by the differential expression of *cadm2a* in response to adipose tissue energy status, leading to changes in cell adhesion dynamics. These changes could activate Rho-GTPase enzymes (e.g., *arhgef25b/GEFT*) (Miller et al. 2014), and the Rho-ROCK signaling (Van Unen et al. 2016). Activated Rho-ROCK then regulates P53 (Shi & Wei 2007), an apoptosis factor that can induce *frmd6* expression, as observed in mammalian neural cells (Park et al. 2024). The induction of *frmd6* activates the Hippo pathway in the hypothalamus, potentially delaying the onset of puberty in females compared to males. Although speculative, as this proposed regulatory axis is based on observations in mammalian cells, it is worthy of future testing in fishes and other vertebrates.

### Sex-specific links between the Hippo pathway and lipid, hormone, and cell adhesion signals in the brain

Our co-expression analysis in the brain showed that there was at least one sex-specific gene co-expression module (GCM) in each sex i.e. gene coexpression was not preserved in the opposite sex (Fig. 4 and 5). This indicates that the majority of gene expression correlations observed within the turquoise GCM in the female brain and the red GCM in the male brain were absent in the opposite sex. This suggests that the functional relationships between these genes were not maintained in the brains of males and females, respectively. In the female brain, the largest identified GCM (turquoise GCM in Fig. 4) was the least preserved in the male brain. The turquoise GCM consisted of genes encoding components of the Hippo pathway and molecular processes including signals mediated by lipid and steroid hormone as well as cell-cell junction and vitamin D biosynthesis (Fig. 4C). The identification of lipid-mediated signals in the brain is not surprising, as the brain is known to sense somatic energy storage (Anderson et al. 2023). While metabolic control of puberty has been studied for decades, the molecular links between fat storage, the brain, and sexual maturation remain underexplored (Anderson et al. 2023). Recent studies have emphasized the role of lipid-sensing signals (e.g., insulin, mTOR, AMPK) in triggering puberty and fat-induced precocious puberty (Thomas Danielsen et al. 2013; Vazquez et al. 2019; Estienne et al. 2021). Interestingly, these signals are known to interact with the Hippo pathway (Estienne et al. 2021; Poon et al. 2013; Holden et al. 2020). Moreover, excess lipids can directly modulate the Hippo pathway through physical interactions with TEADs (Holden et al. 2020). Thus, our findings in the turquoise GCM indicate that there are sex-specific transcriptional signatures of the Hippo pathway components (e.g., *tead1* and *tead3*) and factors mediating lipid sensing and hormonal signals in the brain.

In the male brain, the red GCM (the least preserved in the female brain) comprised genes encoding components of the Hippo pathway and three other molecular processes involved in hormone-mediated signals, developmental growth, and tight junction assembly (Fig. 5). The factors underlying tight junction assembly in the brain are crucial for the formation and integrity of the blood-brain barrier (BBB) (Mullier et al. 2010; Li et al. 2022), and sex hormones influence the tightness of the BBB (Segarra et al. 2021; Dion-Albert et al. 2022). Importantly, the BBB is vital for brain permeability during puberty, and sex-specific differences in BBB permeability have been demonstrated in mammalian studies (Dion-Albert et al. 2022; Lettieri et al. 2023; Solarz et al. 2021). The Hippo pathway is emerging as a key player in BBB formation, homeostasis, and regeneration through the regulation of tight junction proteins (Gong et al. 2019; Hong et al. 2019; Wei et al. 2023). Therefore, the distinct sex-specific transcriptional signatures of these components observed in the salmon brain suggests potential Hippo pathway-dependent differences in BBB tightness, which may underlie sex differences in BBB permeability and lead to distinct brain responses to maturation-related circulating molecules (e.g., lipids) also in this species.

### Conclusions

This study provides significant molecular evidence linking Hippo pathway to sex specific differences in the brain and adipose tissue, which may explain how male sexual maturation often occurs earlier in Atlantic salmon. Our results suggest that females may have higher adipogenesis and lower energy expenditure (lipolysis capacity) compared to males, likely due to sex-specific interactions between PKA and Hippo signaling pathways. In males, increased expression of lipolysis markers in adipose tissue may result in greater energy release, which is sensed in the brain. We also found Hippo-dependent differences in expression of genes encoding tight junction proteins, potentially contributing to greater brain permeability in males. This increased lipid release and potential changes in the brain permeability may underlie sex differences in central lipid sensing process, influencing sex-specific pubertal timing. However, further detailed functional assessments are necessary to validate these suggested differences.

## Acknowledgements

We thank Jaakko Erkinaro for providing the salmon fish used for this study, and Pooja Singh for guiding us through the WGCNA data analysis. We also thank N. Piavchenko, S. Andrew, O. Andersson, T. Aykanat, Y. Czorlich, A. House, M. Lindqvist, N. Lorenzen, O. Mehtälä, K. Mobley, J. Moustakas-Verho, O. Ovaskainen, S. Papakostas, N. Parre, A. Ruokolainen, V. Pritchard, K. Salminen, M. Sinclair-Waters, S. Tillanen, and K. Zueva for help related to gamete stripping, sample processing, tagging, genotyping, phenotyping or fish husbandry, and the staff at the Natural Resources Institute Finland (Luke) hatchery in Taivalkoski hatchery for help during spawning.

## Author Contributions

EPA, CRP, JPV, JK and PVD conceived the study; JPV, PVD and CRP reared and sampled the fish; CRP provided resources; AR, EPA and JPV performed experiments; EPA, JPV and JK developed methodology and analyzed the data; EPA, JPV and CRP interpreted results of the experiments; EPA, JPV and CRP drafted the manuscript, with EPA having the main contribution, and all authors approved the final version of manuscript.

## Ethical approval

The experiments were approved by the Project Authorisation Board (ELLA) on behalf of the Regional Administrative Agency for Southern Finland (ESAVI) under experimental license ESAVI/2778/2018.

## Data availability

All the gene expression data generated during this study are included in this article as supplementary file.

---

**Supplementary File 1: Expression data and statistical analysis.**

